# Rapid generation of precision preclinical cancer models using regulatable in vivo base editing

**DOI:** 10.1101/2022.08.03.502708

**Authors:** Alyna Katti, Miguel Foronda, Jill Zimmerman, Maria Paz Zafra, Sukanya Goswami, Eric E. Gardner, Bianca J. Diaz, Janelle M Simon, Alexandra Wuest, Wei Luan, Maria Teresa Calvo Fernandez, Anastasia P. Kadina, John A Walker, Kevin Holden, Francisco J. Sánchez Rivera, Scott W. Lowe, Lukas E. Dow

## Abstract

Single nucleotide variants (SNVs) comprise the majority of cancer-associated genetic changes and can have diverse effects on protein function. Despite a comprehensive catalogue of SNVs across human cancers, little is known about their impact on tumor initiation and progression. To enable the functional interrogation of cancer-associated SNVs, we developed a murine system for temporal and regulatable in vivo cytosine base editing (iBE). The iBE mice show robust, doxycycline-dependent expression across a broad range of tissues with no evidence of DNA or RNA off-target effects. Transient iBE induction drives efficient creation of individual or multiple SNVs in intestinal, lung, and pancreatic organoids, while temporal iBE regulation allows controlled sequential genome editing. Moreover, in situ delivery of plasmid-based or synthetic sgRNAs to target tissues facilitates the simple and rapid generation of pre-clinical cancer models. Overall, iBE is a powerful in vivo platform to define and interrogate the genetic drivers of cancer.

## INTRODUCTION

Missense and nonsense mutations represent the vast majority of disease-associated genetic changes^1,2^. Cancer cells often harbor thousands of single base pair substitutions^3^ and understanding the impact of specific variants is critical for defining disease drivers and highlighting therapeutic vulnerabilities. While many model systems rely on gene ‘knockout’ or overexpression studies to interrogate the role of specific genes in disease states, ample evidence suggests that individual single nucleotide variants (SNVs), even within the same gene^4–7^ or codon^8,9^, can dictate unique cancer phenotypes and response to targeted therapies.

Mice and organoids are powerful pre-clinical model systems yet engineering cancer-associated SNVs in these complex settings is still laborious and inefficient. Cytosine and adenine base editing (CBE and ABE, respectively) offer the most efficient approach to create targeted (C:G to T:A or A:T to G:C) SNVs^10–13^; however, efficient in vivo BE requires robust expression of editing enzymes that is limited by adequate delivery and can induce antigen-driven immune responses^8,14–21^. Improving the ease, efficiency, and control with which BE and SNVs can be generated in complex cell systems will streamline the functional annotation of disease-associated genetic changes.

Here, we describe a new mouse model carrying an inducible and regulated optimized cytosine base editor (BE3RA^8^) to enable temporal regulation of BE in a wide variety of murine tissues.

We show that inducible and transient expression of a single integrated BE enzyme (iBE) is capable of driving highly efficient base editing in pancreatic, lung, and intestinal organoids without detectable RNA off target effects. Further, we demonstrate that iBE can be used in combination with somatic sgRNA delivery, to build in vivo pre-clinical models of hepatocellular carcinoma (HCC) and pancreas ductal adenocarcinoma (PDAC) harboring specific cancer-associated single nucleotide variants. In all, our study provides a unique tool for efficient modeling of SNVs in physiologically accurate preclinical models to define and test their impact in tumor initiation and progression.

## RESULTS

### Tightly regulatable in vivo expression of an inducible cytosine base editor

To derive mice carrying an inducible base editing (iBE) allele, we injected KH2 mouse embryonic stem cells (ESCs) harboring an expression optimized *TRE-BE3RA* transgene downstream of the *Col1a1* locus^8^ (Figure 1a) into albino B6 blastocysts. High-chimerism (agouti) founders were then backcrossed to C57Bl/6 mice for at least 4 generations before analysis. In the absence of doxycycline (dox), the iBE allele transmitted at normal Mendelian ratios and could be maintained as a heterozygous or homozygous colony (Figure 1b, Supplementary Table 1). We have previously shown that induction of doxycycline-regulated transgenes at the *Col1a1* locus with a constitutively expressed third-generation reverse tet-transactivator (*CAGs-rtTA3*) allele drives uniform expression across a broad range of murine cell types, particularly epithelial tissues^22,23^. To evaluate expression in the iBE mouse, we generated *CAGs-rtTA3;iBE^+/-^* (iBE^het^) mice and treated with dox chow (200 mg/kg) for one week. All tissues examined showed dox-dependent induction of BE3RA that could be reversed following dox withdrawal (dox switched, or SW) (Figure 1c). No tissues showed evidence of sustained BE expression in the absence of dox (Figure 1c). Unlike the uniform induction seen in *GFP-shRNA* transgenes at the *Col1a1* locus^22,23^, BE3RA appeared heterogeneous across multiple tissues (Figure 1d), likely due to stochastic silencing during embryogenesis. We reasoned that presence of a second iBE allele (*CAGs-rtTA3;iBE3^+/+^*, or iBE^hom^) would limit the chance of heritable silencing. Indeed, iBE^hom^ mice showed near ubiquitous expression in liver, pancreas, small intestine, and colon (Figure 1d).

**Figure 1.**
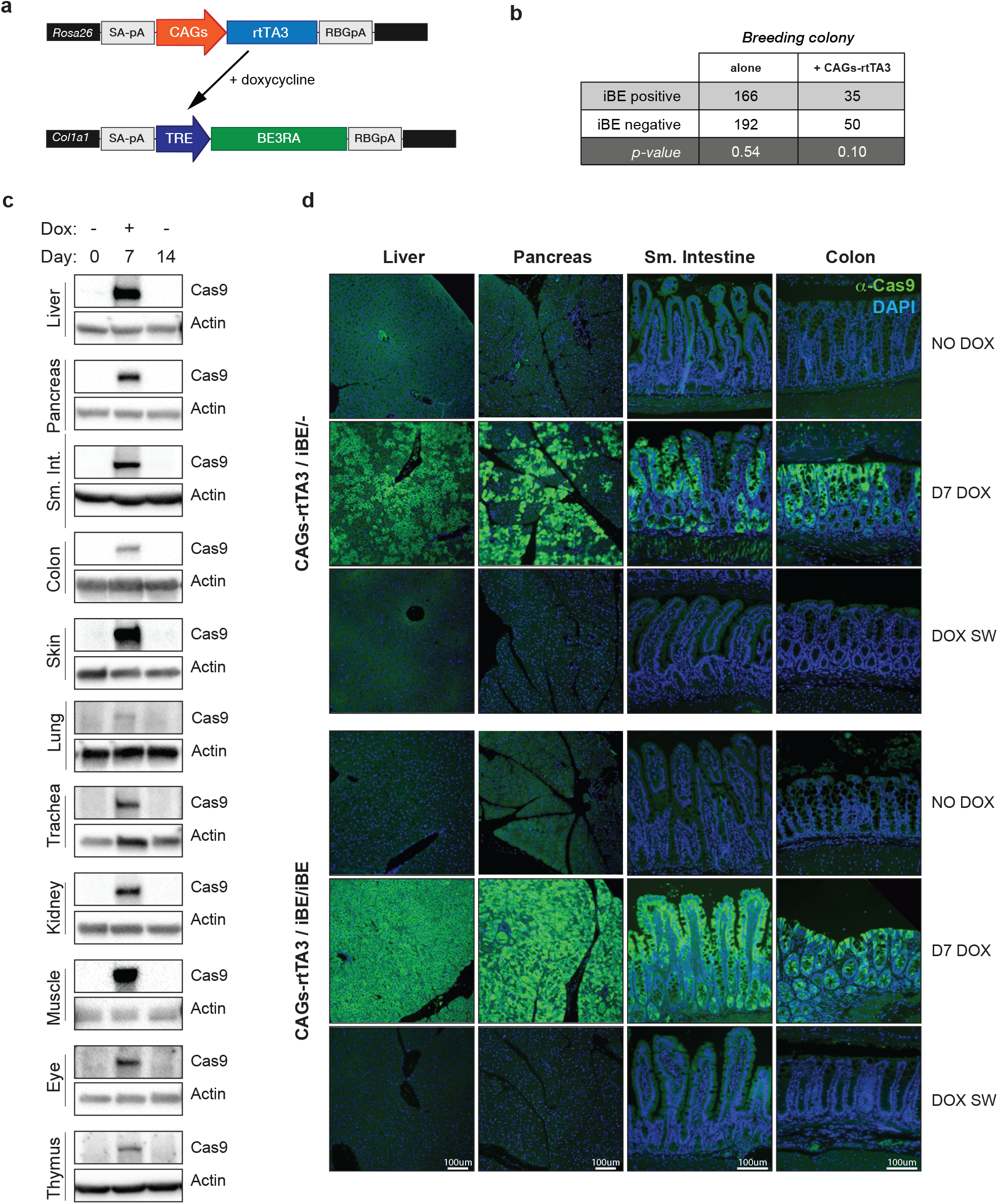
Regulatable BE expression across tissues. **a.** Schematic of breeding scheme for iBE mice containing *R26-CAGs-rtTA3* allele and *Col1a1-BE3RA* allele (targeted knock-in generated in mESCs). **b.** Mendelian transmission of *Col1a1*-targeted iBE knock-in (with and without *R26-CAGs-rtTA3* allele) and associated p-value relative to expected Mendelian inheritance. **c.** Cas9 immunoblot for iBE^het^ mice maintained on normal chow (Day 0, -dox), doxycycline chow (200mg/kg) for 7 days (Day7, + dox), or dox switched from doxycycline chow for 7 days to normal chow for 7 days (D14, -dox) across tissues bulk harvested for protein. ß-actin, loading control. **d.** Immunofluorescent detection of Cas9 protein (CST rabbit α-Cas9 rabbit clone E7M1H #19526) in iBE^het^ (top) or iBE^hom^ (bottom) mice maintained on normal chow (No dox) or doxycycline chow (200mg/kg) for 7 days (Day 7 Dox). Cas9 protein(green), DAPI staining for nuclei (blue) across four tissues analyzed: liver, pancreas, small intestine and colon (left to right).

### iBE enables precise somatic editing in the liver

We and others have previously used transfection or viral-based delivery of base editors to generate somatic mutations in mouse hepatocytes, in situ ^8,14–21,24,25^. Though remarkably effective in liver, somatic delivery of Cas9-based enzymes can result in antigen-mediated immune responses^26–28^. As a first step to measure whether in vivo somatic editing with the iBE allele could promote tumor development, we used hydrodynamic tail vein injection (HTVI) to introduce a *Myc* cDNA in a sleeping beauty (SB) cassette (*SB-Myc*) as well as sgRNAs targeting *Apc*, *Ctnnb1*, or *Trp53* ^8,29,30^ designed to engineer known cancer-linked SNVs. To drive BE enzyme expression, iBE mice were maintained on dox for one week prior to injection and one-week post-injection (Figure 2a). Six weeks following HTVI, tumors were palpable, and bulk tumor tissue was harvested for sequencing and histological analysis (Figure 2a). Most mice (12/13) injected with *SB-Myc* and a control sgRNA targeting a non-genic region (CR8) had no macroscopically visible tumors, but showed small, well-circumscribed regions on histological sections, which were also observed in *SB-Myc* only animals (Figure 2b). Consistent with an established role for WNT in a subset of HCC ^8,29,30^, *SB-Myc;Apc^Q1405X^* and *SB-Myc;Ctnnb1^S33F^* mice showed markedly enhanced tumor growth (Figure 2b).

**Figure 2.**
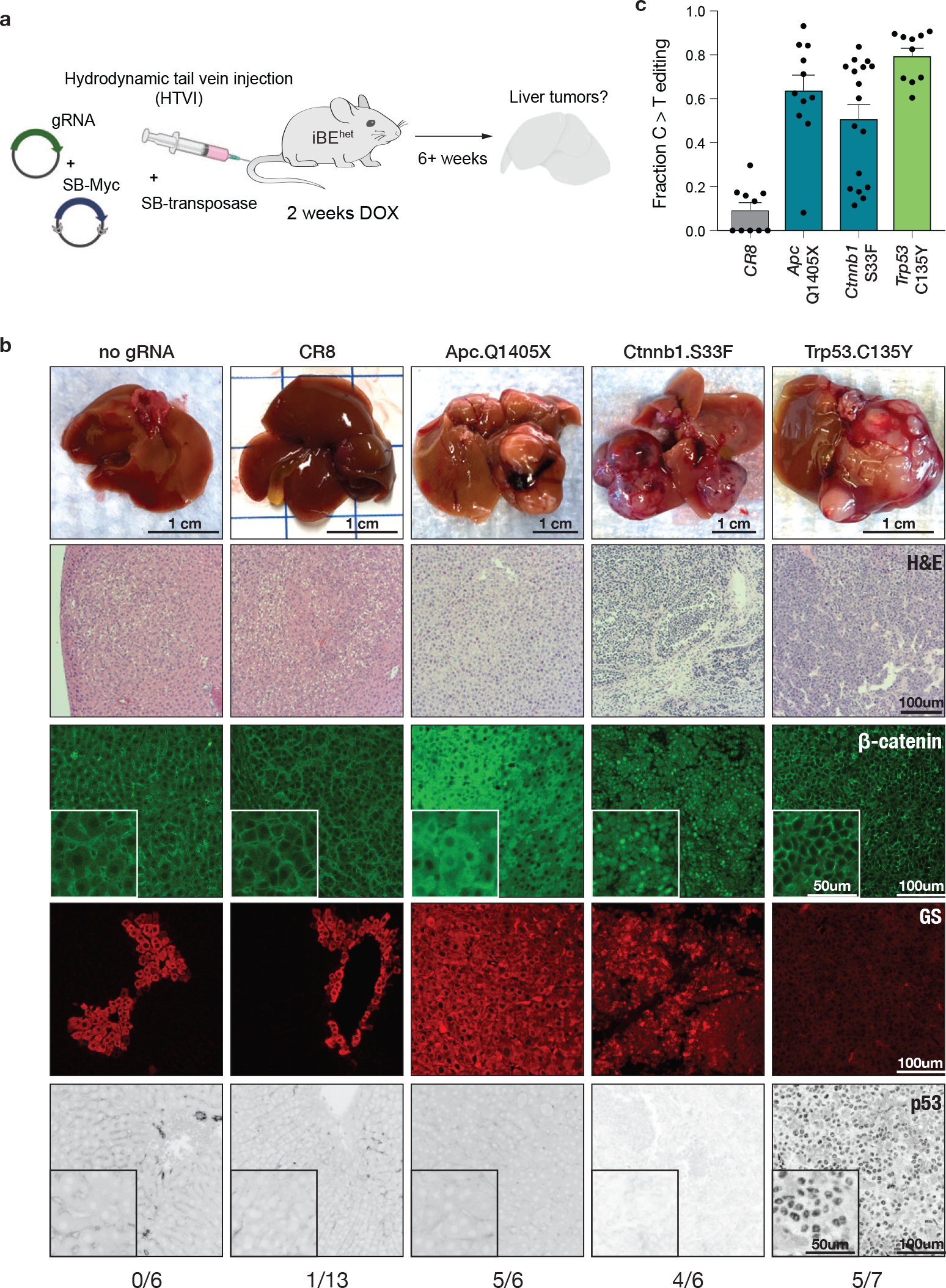
In situ base editing with iBE drives liver tumors. **a.** Schematic for experimental setup of hydrodynamic teil vein (HTVI) injection mediated delivery of plasmid gRNA and sleeping beauty transposon mediated integration of cMyc cDNA (SB-Myc) in the liver of iBE mice maintained on dox for 1 week surrounding injection. Post injection, mice are monitored for tumor development and palpable tumors are harvested for tumor histological and sequencing analysis. **b.** Brightfield images of liver after harvest targeted according to the experimental pipeline in a) and with the corresponding gRNA listed (top). Hematoxylin and eosin (H&E) staining (10X, 2nd row) of corresponding livers. Immunofluorescent staining of total ß catenin (green, 3rd row), glutamine synthetase (GS, red, 5th row). Fraction of number of mice with palpable tumors over number of mice injected is below each gRNA column. **c.** Targeted deep sequencing analysis of target C:G to T:A conversion in tumors collected in b) delineated by gRNA/target site. Each point corresponds to an isolated bulk tumor. (n=3 mice minimum for a given gRNA target).

Targeted amplicon sequencing revealed a high proportion of the expected SNVs, with low rates of insertions or deletions (indels). Absolute editing rates in bulk tumor tissue were variable, likely due to the presence of admixed stroma and immune cells (Figure 2c and Supplementary Figure 1a-c). Notably, *SB-Myc;Ctnnb1^S33F^* tumors showed lower overall editing rates that Apc^Q1405X^ mutant tumors, consistent with the notion that heterozygous *Ctnnb1^S33F^* mutations are sufficient to activate WNT signaling, while Apc requires inactivation of both alleles. Indeed, both Apc^Q1405X^ and Ctnnb1^S33F^ tumors showed accumulation and mislocalization of β-catenin protein and elevated expression of glutamine synthetase (GS), a WNT target that is normally restricted to pericentral hepatocytes surrounding the central vein (Figure 2b). Consistent with a strong tumor suppressive role for p53 in HCC, introduction of an sgRNA targeting *Trp53* (Trp53^C135Y^) accelerated tumor growth, with 5/7 mice showing multi-focal tumors, with high levels of on-target editing (Figure 2b,c and Supplementary Figure 1d). Similar to previously characterized p53 hotspot mutations^31–34^, C135Y resulted in p53 protein stabilization and nuclear localization (Figure 2b,c and Supplementary Figure 1d). These data show that the iBE mouse enables temporally regulated target editing and can be used to generate in vivo liver cancer models through controlled and precise induction of cancer-relevant SNVs.

### Efficient multiplexed editing in iBE organoids

Organotypic cell culture models or ‘organoids’ are a powerful system to study epithelial biology. We and others have used organoids to reveal the contribution of cancer-associated nonsense and missense mutations for cell behavior and drug response^9,35,36^; however, efficient introduction of such mutations using base editing is a substantial practical challenge in scaling up the generation of large collections of tailored model systems. To determine if cells derived from iBE mice could streamline the creation of targeted mutations ex vivo, we generated organoids from small intestine, pancreas, and basal cells from the trachea of iBE^het^ mice. Each culture showed robust inducible and reversible expression of BE3RA (Supplementary Figure 2a), and organoids transduced with BE activatable GFP^GO^ reporter^37^ showed editing efficiencies ranging from 40-90% (Figure 3a, Supplementary Figure 2b,c).

**Figure 3.**
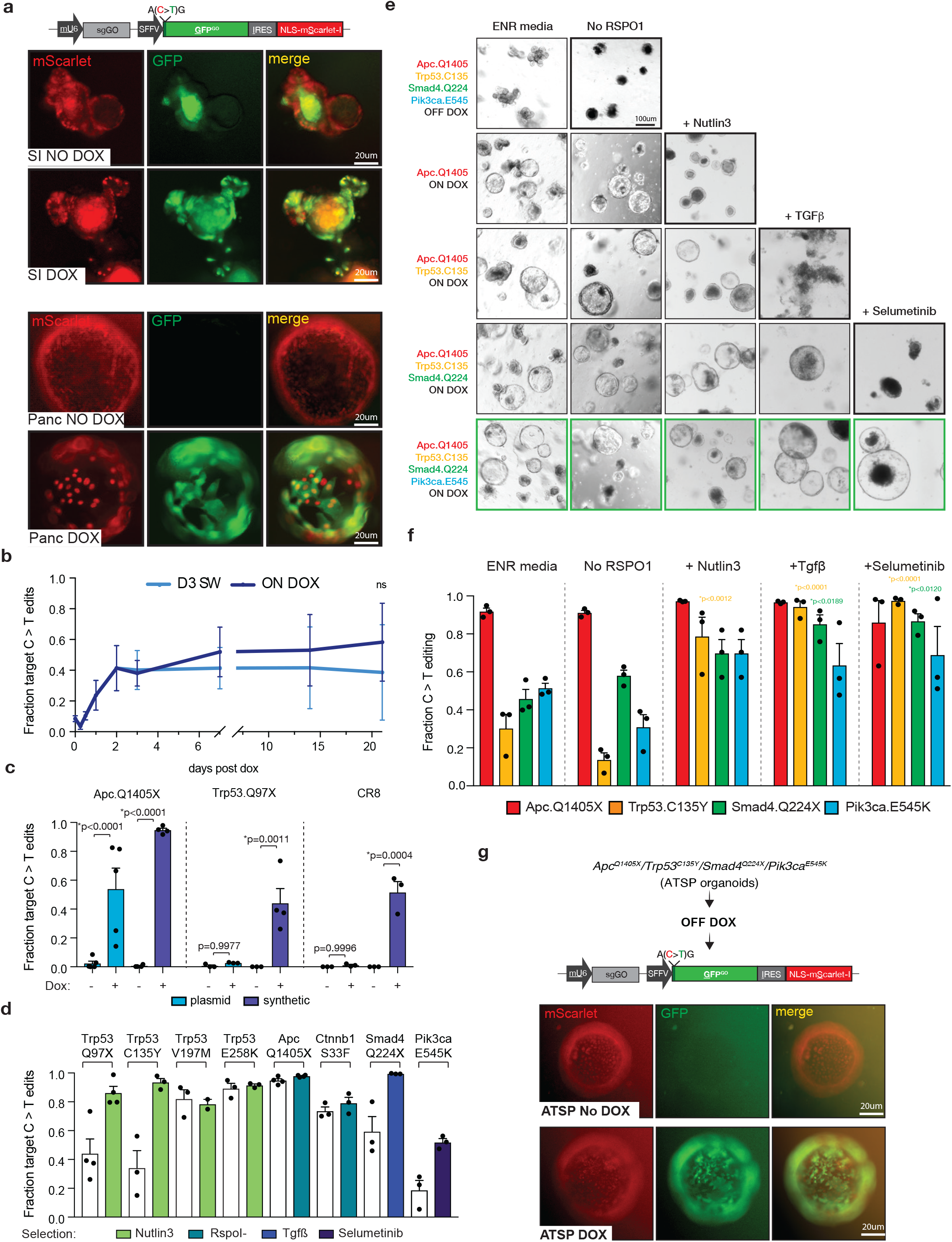
Efficient base editing in ex vivo derived iBE organoids. **a.** Fluorescence-based imaging of small intestinal (top) and pancreatic (bottom) organoids containing stable integration of GFPGO lentiviral construct (schematic for reporter above images) cultured without (no dox) or with dox. Constitutive mScarlet marker for the reporter is shown in red (left column), base editing activatable GFP is shown in green (middle column), Merged images are depicted on the right. **b.** Targeted deep sequencing quantification of target C:G to T:A conversion at the Apc^Q1405X^ locus in 2D small intestinal derived iBE cell line (constitutive *shApc;shTrp53;Kras^G12D^* cDNA) with lentiviral integration of a gRNA (LRT2B) targeting Apc^Q1405X^ at time points after dox is added to the culture media (0 to 15 days) (dark blue),or dox was cultured in the media briefly for 3 days (light blue). **c.** Targeted deep sequencing quantification of corresponding target C:G to T:A conversion in small intestinal iBE organoids nucleofected with various plasmid (light blue) or synthetic (indigo) gRNAs (Apc^Q1405X^, Trp53^Q97X^, CR8^OS2^) with and without dox conditions for 2 days surrounding nucleofection. **d.** Targeted deep sequencing quantification of target C:G to T:A conversion in small intestinal iBE organoids nucleofected with various synthetic gRNAs targeting cancer associated SNVs cultured in dox or 2 days surrounding nucleofection, and either unselected (-) or selected with corresponding functional selective media condition (+: Trp53^mut^, nutlin selection; Smad4^mut^, Tgfß selection; Pik3ca^ACT^, Selumetinib selection; Ctnnb1^ACT^, RspoI withdrawal). **e**. Brightfield images of small intestinal iBE organoids targeted with various gRNA combinations and dox conditions (left) taken through sequential selection through Rspo1 withdrawal, Nutlin selection; Tgfß selection, and Selumetinib selection. Bolded black boxes are conditions failing to survive selection. Bolded green boxes are the quadruple targeted organoids (with dox) surviving all four selection conditions. **f.** Targeted deep sequencing quantification of target C:G to T:A conversion in small intestinal iBE organoids nucleofected with 4 synthetic gRNAs in e (green boxes) at each gRNA target loci (Apc^Q1405X^, Trp53^C135Y^, Smad4^Q224X^, Pik3ca^E545K^. Media conditions and corresponding organoid genotype and sequencing information is grouped and listed above. **g.** Schematic describing BE targets and selective culture conditions of organoids prior to GFP^GO^ infection and fluorescence-based imaging of small intestinal iBE^hom^ organoids from (**f**) infected with GFP^GO^ and cultured off dox or on dox. (organoids were pulsed with dox to make the original four endogenous mutations and are pulsed a second time to see induction of BE)

To closely assess editing dynamics, we generated an immortalized 2D cell line from small intestinal organoids and transduced cells with *LRT2B*^8^ expressing the *Apc^Q1405X^* sgRNA and measured target editing over 3 weeks and following withdrawal of dox at 72hrs. Target editing was detectable as early as 12h after dox addition, reaching 80% of total BE within 48hrs. (Figure 3b). Overall editing increased only moderately beyond day 3, whereas dox withdrawal at this time prevented further editing at the target site (Figure 3b). Consistent with previous experience with this sgRNA, indels were rare at D3 (~1% total reads) but increased 4-fold when maintained on dox over 3 weeks (Supplementary Figure 3d).

We next asked whether iBE could improve the efficiency of building complex organoid-based models of cancer. To limit the introduction of exogenous and potentially immunogenic components we opted for transient transfection of sgRNAs by nucleofection. Organoids were cultured in dox media for 2 days before and after nucleofection (4 days total) to transiently induce the editor and align BE protein expression with sgRNA expression; editing was quantified 7d post-transfection by amplicon sequencing. Nucleofection of the Apc^Q1405X^ sgRNA in a U6 expression plasmid (*LRT2B*) induced ~50% target editing, while *Trp53^Q97X^* and *CR8^OS2^* gRNAs showed C>T editing below 5% (Figure 3c). In contrast to plasmid-based delivery, nucleofection of dox-treated organoids with chemically stabilized synthetic sgRNAs led to significantly higher editing, up to 43-fold higher in the case of CR8^OS2^ (Figure 3c, Supplementary Figure 2e; p = 0.0005). Using the more efficient synthetic sgRNA approach, we next tested a range of additional BE sgRNAs predicted by BE-SCAN (https://dowlab.shinyapps.io/BEscan/)^38^. In unselected populations, 7d post-transfection, C>T editing efficiencies ranged from ~20%-90% (Figure 3d). Functional selection for mutant organoids (RSPO withdrawal for Ctnnb1^S33F^, or TGFβ for Smad4^Q224X^, Nutlin3 treatment for p53 mutations, and Selumetinib treatment for Pik3ca^E545K^) enriched C>T editing 80-95% for tumor suppressor (Smad4 and Trp53) and 50-80% for oncogenes (Pik3ca and Ctnnb1) (Figure 3d, Supplementary Figure 2f). Given the high efficiency of engineering individual target mutations, we next asked whether iBE organoids could be used for rapid multiplexed editing. Using iBEhom organoids to maximize the likelihood of efficient enzyme expression in all cells, we delivered a combination of four sgRNAs - *Apc^Q1405X^*, *Trp53^C135Y^*, *Smad4^Q224X^*, and *Pik3ca^E545K^* - to model frequently observed colon cancer mutations; notably, three of the four sgRNAs used show moderate editing efficiency (20-45%), thereby providing a test of multiplexed editing in non-ideal circumstances. Target editing for each site in the bulk, unselected populations resembled that seen with individual transfections in iBE^het^ cells (Figure 3e, Supplementary Figure 2g). Iterative functional selection for each mutation resulted in an organoid population with ~80-90% C>T editing for *Apc*, *Trp53*, and *Smad4* (Figure 3f), while *Pik3ca* editing reached ~65%, consistent with the notion that heterozygous mutations are sufficient to activate PI3K signaling. Control organoids receiving all 4 sgRNAs in the absence of dox did not survive functional selection (Figure 3e). Together, these data show that iBE provides an efficient system for engineering disease relevant SNVs and can be used to quickly generate genetically complex cancer models.

### Undetectable off-target editing in iBE organoids

Previous studies have reported that BE enzymes can produce widespread, sgRNA-independent off-target RNA editing^39^. To determine if RNA editing could be a concern in iBE mice, we analyzed mRNA from pancreatic iBE organoids carrying the GO reporter (Supplementary Figure 3a). iBE pancreatic organoids transduced with GFP^GO^ (OFF dox) were treated with dox for 4 days (D4), 8 days (D8), or treated for 4 days and switched off dox for 4 days (D4 SW). As previously shown (Figure 3a, Supplementary Figure 3b,c), dox treatment led to GFP expression in ~60% of cells (Supplementary Figure 3b). Analysis of mRNA from OFF, D8, and D4 SW transcriptomes revealed that any variance between samples was predominantly driven by differences between biological (mouse) replicates rather than BE expression (Supplementary Figure 3c, Supplementary Table 2). In contrast with a large number of transcriptional changes observed in BE3 transfected HEK293T cells^39^ (Supplementary Figure 3d), dox-treated and dox-switched iBE organoids showed few differentially expressed genes compared to OFF dox cultures, with the largest expression change being the BE enzyme itself (Supplementary Figure 3e, Supplementary Table 2). Similarly, we did not detect any increase in the level of C>U edited RNA transcripts following BE3RA expression (Supplementary Figure 3g). One possible explanation for this difference is relative levels of enzyme expression between experimental systems. BE3 transcript was ~6000-fold higher in transfected HEK293T cells than dox-treated iBE cells (Supplementary Figure 3f), suggesting that reduced BE enzyme provides effective DNA editing while avoiding off-target RNA effects. We further assessed the possibility of potential DNA off-target effects using iBE targeted ESCs expressing Apc^Q1405X^ and Pik3ca^E545K^ sgRNAs. The use of ESCs enabled the reliable growth and isolation of clonal populations following iBE induction by dox treatment. In total, we sequenced 60 (non-dox) control and 60 dox-treated clones (in pools of 10) using ultra deep sequencing (800-1000x coverage) of a focused panel of cancer-relevant genes (MSK-IMPACT^40^). We saw no evidence of increased C>T DNA editing in dox treated cells relative to controls (Supplementary Figure 4, Supplementary Table 3). In all, these data highlight that transient and controlled induction of the iBE transgene enables precise C>T target editing without any detectable widespread DNA or RNA off-target effects.

### Engineering oncogenic missense mutations in the pancreas with iBE

The generation of *in vivo* cancer models harboring recurrent cancer-associated missense mutations is a critical step in understanding their role in carcinogenesis. To test the potential for using iBE in the generation of in situ cancer models, we used an electroporation-based approach^41,42^ to introduce a targeted Trp53 mutation into the mouse pancreas alongside a SB cassette containing *Kras^G12D^* (*SB-Kras*) (Figure 4a). iBE^het^ mice targeted with Trp53 gRNAs showed rare incidence of tumor development (1/10 mice, not shown); however, iBE^hom^ mice showed induction of large pancreatic tumors in 4/7 mice by 5-8 weeks (Figure 4b, Supplementary Figure 5). Similar to genetically engineered *Kras;p53* driven Cre models (KPC)^31^, tumors contained CK19+ ductal islands with substantial surrounding stroma expressing alpha-smooth muscle actin (α-SMA, Figure 4b). Sequencing of bulk tumors revealed precise on target C>T editing with minimal indels; absolute editing percentages were low, likely due to abundant non-tumor cells in bulk sample, as frequently observed in human and mouse PDAC (Figure 4c).

**Figure 4.**
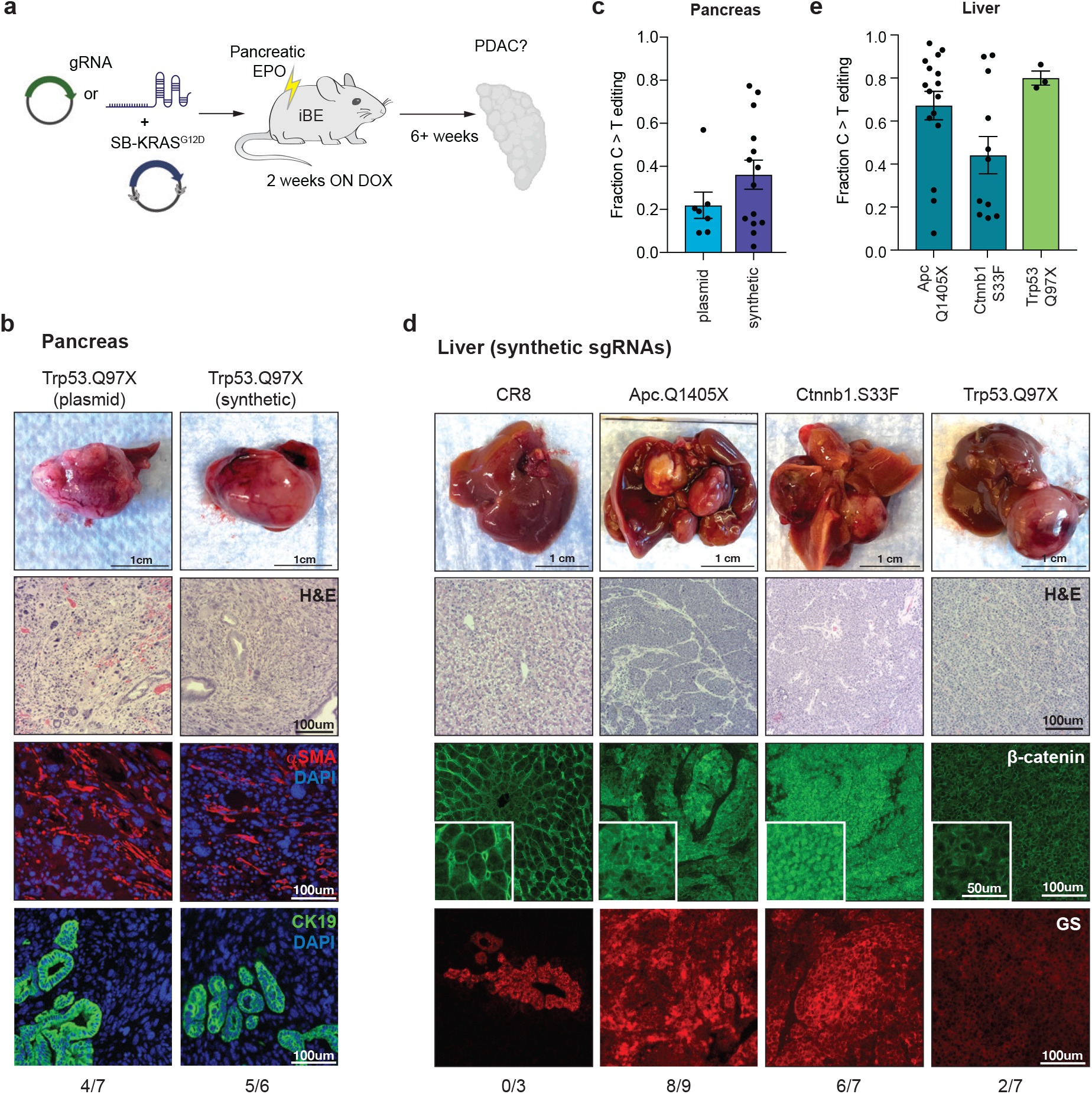
Efficient engineering of missense mutations in pancreatic tumor models. **a.** Schematic for experimental setup of pancreatic electroporation mediated delivery of gRNA and sleeping beauty transposon mediated integration of *Kras^G12D^* cDNA (*SB-Kras*) in the pancreas of iBE^hom^ mice maintained on dox for 2 weeks surrounding electroporation. Post electroporation, mice are monitored for tumor development and palpable tumors are harvested for tumor histological and sequencing analysis. **b.** Brightfield images of pancreas tumor with spleen attached (top row) using plasmid or synthetic gRNA targeting Trp53^Q97X^. Hematoxylin and eosin (H&E) staining (10X, 2nd row) of pancreatic tumors electroporated as in panel (**a**) for gRNAs listed. Immunofluorescent staining of alpha smooth muscle actin (αSMA, red, 3rd row) and cytokeratin-19 (CK19, green, 4th row) counterstained with DAPI (blue). Fraction of number of mice with palpable tumors over number of mice injected is below each gRNA column. **c.** Targeted deep sequencing analysis of target C:G to T:A conversion in tumors collected in (**b**) for plasmid gRNA (left) and synthetic gRNA (right). Each point corresponds to an isolated bulk tumor. (n=3 mice minimum for a given gRNA target). **d.** HTVI delivery of synthetic gRNAs with *SB-Myc* as in Figure 2. BF, H&E images, and IF staining for ß-catenin (green) and glutamine synthetase (GS, red) in livers with tumors. **e.** Quantification of target C:G to T:A conversion in tumors found below.

In an effort to improve the efficiency of developing in vivo models with iBE and eliminate the need for sgRNA cloning, we tested the delivery of chemically-stabilized but non-encapsulated (‘naked’) synthetic sgRNAs. Electroporation of the pancreas with *SB-Kras^G12D^* and synthetic *Trp53^Q97X^* sgRNAs in iBE^hom^ mice showed highly penetrant tumor growth (5/6 mice), efficient C>T editing and identical histology to what was observed with plasmid-based delivery (Figure 4c). Likewise, synthetic sgRNAs targeting *Apc*, *Ctnnb1* or *Trp53* coupled with *SB-Myc* drove consistent tumor formation in the liver following HTVI (8/9, 2/7, and 6/7 mice, respectively)(Figure 4d). For both nonsense (Apc^Q1405X^ and Trp53^Q97X^) and missense (Ctnnb1^S33F^) mutations, we saw high on-target C>T editing and low indel formation (Figure 4d). Thus, iBE mice provide a platform for rapid and easy generation of disease-associated SNVs in multiple organ sites *in situ*.

## DISCUSSION

The generation of model systems that faithfully recapitulate the genetic alterations observed in human disease is a key step in developing precision treatment strategies. Here, we set out to produce an efficient and regulated platform to streamline the creation of such pre-clinical disease models. The iBE platform enables efficient creation of targeted nonsense and missense mutations in vivo in somatic tissues, and in cells and organoids derived from mice. Further, the system supports multiplexed and/ or sequential editing with synthetic sgRNAs, thus providing a rapid approach to engineer complex genetic combinations often seen in human cancers.

Previous work has demonstrated the potential of in vivo BE for engineering SNVs^8,14–21,24,25^, though these approaches have relied on exogenous delivery of BE enzymes using transfection, split inteins or single editors in AAV vectors, or engineered virus like particles (eVLPs). These approaches can catalyze highly efficient editing in a subset of tissues (i.e. the liver, muscle, brain, and eye)^8,14–21,24,25^, though may suffer from unintended immune recognition of Cas9-derived antigens^26–28^. iBE broadens the number of tissues that can be targeted with CBEs for disease modeling and as it is encoded in the genome, may avoid complications of immunogenicity when induced in somatic tissues. Annunziato et al previously described a genomically-encoded Cre-activatable BE mouse using the pre-optimized BE3 enzyme^43^. This transgenic mouse demonstrated the ability to induce target editing in the mammary gland and drive tumor development in combination with Myc; however, perhaps due to the sustained expression of BE3 from a constitutive promoter, target sites often accumulated indels rather than desired SNVs^43^. In the present study, we saw minimal indel formation across 10 different target sites both in vitro and in vivo. Yet, as expected, we saw a modest accumulation of indels in cells maintained on dox over 3 weeks in culture. Thus, the reversible induction of enzyme expression in iBE provides a key improvement that limits indel formation at endogenous genomic targets.

Our goal was to develop a system for creating precise, genetically-defined pre-clinical models. As such, we thoroughly explored sequence-independent DNA and RNA off-target effects following dox induction in iBE cells. Rather than performing whole genome sequencing on a small number of post-editing clones, we opted for targeted deep sequencing of cancer-relevant genes contained in the MSK-IMPACT panel^40^. While this approach does not measure editing across the entire genome, it enabled the analysis of 120 individual ESC clones. Surprisingly, we observed a higher total number of SNVs in dox-treated clones, but there was no increase in APOBEC-mediated (C>T) mutation signature. These data suggest that iBE does not induce widespread sequence-independent DNA off-targets in cancer-associated genes, but we cannot rule out the possibility of rare mutations elsewhere in the genome. Similarly, but in contrast to published data^39,44^, we saw no evidence of sequence-independent off-target RNA editing. It is possible this difference is due to changes in the experimental conditions of our work, although it is also likely that the relatively modest level of enzyme expression from the single copy transgene can maintain on-target DNA editing while minimizing potential off-target effects (Supplementary Figure 3), thus providing an optimized platform for preclinical modeling using BE.

In all, the iBE mouse simplifies base editing in complex systems and provides an easy to use and reliable method for engineering cytosine base edits for disease modeling Our proof-of-concept studies demonstrate the utility of this platform for ex vivo and in vivo target editing. Further, given the broad expression of iBE across all tested tissues, and the ability to control spatial BE activity via Cre-dependent or tissue restricted rtTA alleles, our model is a powerful tool to engineer and study missense mutation in vivo for many disease applications.

## Acknowledgements

We thank members of the Dow lab for advice and comments on preparation of the manuscript. The work was supported by a project grant from the National Institutes of Health (NIH/NCI; R01CA229773), P01 CA087497 (SWL), a MSKCC Functional Genomics Initiative (FGI) grant (SWL), and an Agilent Technologies Thought Leader Award (SWL), and support from Synthego Inc. under a Synthego Innovator Award (LED). A.K. was supported by an F31 Ruth L. Kirschstein Predoctoral Individual National Research Service Award (F31-CA247351-02). B.J.D. was supported by an F31 Ruth L. Kirschstein Predoctoral Individual National Research Service Award (F31-CA261061-01). E.E.G. is the Kenneth G. and Elaine A. Langone Fellow of the Damon Runyon Cancer Research Foundation (DRG-2343-18). FJSR was supported by the MSKCC TROT program (5T32CA160001), a GMTEC Postdoctoral Researcher Innovation Grant, and is an HHMI Hanna Gray Fellow. SWL is an HHMI investigator. The content is solely the responsibility of the authors and does not necessarily represent the official views of the NIH.

## Authors’ contributions

A.K. designed and performed experiments, analyzed data and wrote the paper. M.F., J.Z., M.P.Z., S.G., E.G., J.S. and W.L, B.J.D., M.C.F., K.H., and F.J.S-R., performed experiments and/or analyzed data. S.W.L. supervised experimental work. L.E.D designed and supervised experiments, analyzed data and wrote the paper.

## Methods

### Animals

All animal experiments were approved by the Weill Cornell Medicine Institutional Animal Care and Use Committee (IACUC) under protocol 2014-0038 or by the MSKCC Institutional Animal Care and Use Committee under protocol 11-06-018. ES cell-derived mice were produced by injection into albino B6 blastocyst by the transgenic targeting core facility at NYU School of Medicine. High chimera (agouti) founders were backcrossed to C57Bl/6 mice for at least 4 generations before analysis. iBE^het^ mice were generated through breeding with C57Bl/6 mice containing a R26-CAGs-rtTA3 allele (Supplementary Table 1). iBE^hom^ mice were generated through breeding iBEhet progeny. Mice were genotyped by Col1a145, R26, and CAGs-rtTA3 PCRs using EconoTaq PLUS (Lucigen #30033-2). Doxycycline chow (food pellets) were administered for 1 or 2 weeks (as specified) at 200mg/kg (Envigo #TD.180625). Mice were manipulated experimentally (organoids, injection, or electroporation) at 8-12 weeks of age. Male and female mice were used for all studies.

### Cloning

All plasmid sgRNAs were cloned into the BsmBI site of LRT2B^8^. Oligos for gRNA cloning are found in Supplementary Table 4.

### ES cell targeting

KH2 mouse embryonic stem cells (ESCs) harboring a TRE-BE3RA transgene at the Col1a1 locus were engineered as previously described^8^. Briefly, the BE3RA cDNA from Lenti-BE3RA was cloned into the Col1a1 targeting vector containing a TRE-promoter element (cT)^45^.

### Cells

HEK293T (ATCC CRL-3216) cells were purchased from the ATCC. Stocks were tested for mycoplasma routinely every 6 months and maintained in Dulbecco’s Modified Eagle’s Medium (DMEM, Corning # 10-013-CV) containing 1% Pen/Strep (Corning #30-002-Cl) and 10% FBS at 37C with 5% CO2.

### Organoid culture and transduction

Murine small intestine organoids from the indicated genotypes were isolated and maintained as previously described^46^. Isolation of murine pancreatic ductal organoids was done modifying previously described protocol^35,47^. Isolation of murine pulmonary basal cell spheroids was performed using tracheas pooled from 3 animals. Animals were euthanized by inhaled carbon dioxide, sprayed down with 70% ethanol and then sheathed. Following gross dissection of the thoracic cavity, animals were cardiac perfused with PBS through the left ventricle and tracheas were cut away from the bronchial tree, capping at the submucosal glands. Single cell suspensions were generated using a gentleMACS Octo Dissociator and a mouse lung dissociation kit (Miltenyi #130-095-927) on the m_lung_02 protocol. Crude suspensions were then passed through 70um mesh filters and rinsed with 10cc of cold FACS buffer (PBS + 2% FBS + 2mM EDTA). Cells were pelleted at 500g for 3min, red blood cells lysed for 3min using 5cc of ACK lysis buffer (Thermo Fisher #A1049201) and then quenched with 20cc of FACS buffer. Cell pellets were resuspended in FACS buffer, filtered through 70um cell strainer FACS tubes and counted (Nexcelom Cellometer Auto X4). Cd31/Cd45 cells were depleted using 10uL each of Cd31 (Miltenyi #130-097-418) and Cd45 (Miltenyi #130-052-301) microbeads per 107 total cells and passed through an LD depletion column (Miltenyi #130-042-901). Cells were seeded in a 6.5mm transwell insert at ~20K cells in 200uL of a 50% Matrigel suspension (BD Biosciences, 354230) per transwell. Matrigel/cell mix was incubated for 15 min at 37C to allow for solidification and base media with Primocin^®^ (InvivoGen) was added to top and bottom chambers of transwell. Base media constitutes: DMEM/F12 + HEPES (15mM) + Sodium Bicarbonate (3.6mM) + L-glutamine (4mM)+ Insulin (10ug/mL) +Tranferrin (5ug/mL) (or ITS,1x, Sigma Aldrich #I3146) + Cholera Toxin (0.1ug/mL, Sigma Aldrich C9903) + EGF (25ng/mL)+ Bovine Pituitary Extract (30ug/mL, Sigma Aldrich, P1476)+ FBS (5%) + Retinoic Acid (0.05uM). During the first 48hrs of seeding or passage, 1uM Y-27632 (MedChemExpress; HY-10583) was added to the base media. Media was changed every 2 days and cells were passaged after ~1 week and every ~3-4 days for subsequent passages. Organoids were transduced as previously described.

### Generation of 2D lines from organoids

To engineer immortalized 2D lines, 3D small intestinal organoids were transduced with a lentiviral all-in-one Kras^G12D^ cDNA and MultiMiR tandem knockdown cassette (shApc-shTrp53)^48^. After selection in media without RSPOI and Nutlin3 (10umol/L), organoids were split onto plates coated with Rat Collagen I in PBS (Gibco, #A10483-01, 30ng/mL) for 30 minutes at 37C prior to plating. Cells were passaged on Collagen coated plates 3-5 times and then split to plates without Collagen. 2D cells were transduced with lentivirus as previously described^37^.

### Flow cytometry

Cells were trypsinized and organoids were mechanically dissociated followed by TrypLE treatment at the indicated time point. Dissociated cells were resuspended in 500uL of FACS buffer (PBS/2%FBS/3mM EDTA) with DAPI to stain dead cells. Flow cytometry assays were carried out on a Thermo Fisher 2018 Attune NxT flow cytometer at a flow rate of 500ul/min. At least 25,000 events from the single cell population gating were recorded, and gates set as shown (Supplementary Figure 6). All experiments were performed in replicates from independent mouse lines as annotated.

### Organoid Nucleofection

Three days before nucleofection organoids were split for one well in a 12 well plate per condition and cultured in full media (ENR; 50ng/mL EGF, Invitrogen, and 50nMol LDN-0193189, Selleck Chemicals + RSPO1-conditioned media). Two days prior to nucleofection, media was changed to EN (50ng/mL EGF, Invitrogen, and 50nMol LDN-0193189, Selleck Chemicals) + Y-27632 (10umol/L) + CHIR99021 (5umol/L) and with or without doxycycline as noted (500ng/mL). On day of nucleofection, media is removed, and organoids mechanically dissociated in cold PBS by pipetting (50X). Organoid suspension was pelleted by spinning at 1200rpm for 4 min at 4C and resuspended in 100uL TrypLE (Invitrogen #12604) followed by incubation in bead bath at 37C for 5 mins. ~300uL of cold PBS was added followed by mechanical dissociation of organoids by pipetting (50X), and then washed with cold PBS. Per condition, nucleofection mix was prepared as follows: 16.4 uL Primary P3 Buffer (Lonza kit, #V4XP-3032), 3.6 Supplement 1 (Lonza kit, #V4XP-3032), and 1ug of plasmid DNA or 200pmol of chemically stabilized synthetic RNA (Synthego Corp.). For multiplexing experiments, total gRNA concentrations were kept constant and divided evenly by number of gRNAs in that condition. Pelleted organoids were resuspended in 20uL of nucleofection mix and transferred to electroporation chamber (Lonza kit, #V4XP-3032, 96-well format) for electroporation using Lonza X Unit Nucleofector under the [ES, mouse] protocol. Organoids were recovered in 70uL of media and washed once. Pelleted organoids were plated in original volume of Matrigel (BD Biosciences, 354230) and cultured in EN + Y-27632 + CHIR +/- dox media for 2 days and subsequently replaced with full media or selection conditions.

### Organoid functional selection

To select for WNT activating mutations, exogenous RSPO1 was removed from the media. To select for loss-of-function Trp53 mutations, Nutlin3 (5umol/L) was added to the media and organoids were cultured for 10 days. To select for Smad4 alterations, recombinant TGFB1 (5ng/mL) was added to the media and organoids cultured for 7 days. To select for Pik3ca activating mutations, Selumetinib (1ug/mL) was added to the media and organoids cultured for 14 days. Organoids were split as usual throughout selection conditions.

### Genomic DNA isolation

Cells and organoids were dissociated and pelleted at the indicated time point. Cells were lysed as previously described^8^. Tumor nodules were micro-dissected and homogenized using a 5mm stainless steel bead (Qiagen, #69989) and a Tissue LyserII (Qiagen) in 150uL genomic DNA lysis buffer for 3 mins at a frequency of 30 hz/s and immediately cooled for 5 mins on ice. Tumor suspension was then lysed and gDNA isolated identical to cells^8^.

### PCR amplification for sequencing

Target genomic regions of interest were amplified by PCR using primers in Supplementary Table 2. PCR was performed with Herculase II Fusion DNA polymerase (Agilent Technologies, #600675) according to the manufacturer’s instructions using 200 ng of genomic DNA as a template and under the following PCR conditions: 95C × 5 min, 95C - 0:30; 57C - 0:30; 72C - 0:20 × 39 cycles, 72C × 5 min. PCR products were confirmed using Qiaxel and purified using QIAquick PCR Purification Kit (Qiagen # 28106). PCR products were quantified by NanoDrop (Thermofisher Scientific Inc) and normalized to 20ng/uL in EB buffer. Targeted amplicon library preparation and NGS sequencing (MiSeq; 2 x 250bp) were performed at Azenta (previously GENEWIZ, Inc.) and analyzed using CRISPResso2. Raw MiSeq fastq files have been deposited in the sequence read archive (SRA) under accession PRJNA859154.

### Protein analysis

#### Organoids

A 6-well of organoids was collected in Cell Recovery Solution (Corning, #354253) and incubated on ice for 30min - 2 hours, washed with PBS 3 times to removed residual Matrigel. Organoid pellets were resuspended in 100uL RIPA buffer, and centrifuged at 13,000 rpm at 4C to collect protein supernatant.

#### Tissue

A 2mg piece of each tissue was collected at indicated time points and immediately processed or snap frozen and stored at −80C. Tissue was homogenized in 150uL of RIPA buffer with protease and phosphatase inhibitors by bead homogenizer (Tissue LyserII, Qiagen) for 3 mins at a frequency of 30 hz/s and immediately cooled for 5 mins on ice. The following antibodies were used for western blotting analysis of organoids and tissues: Cas9 (Biolegend, #844301) (1:500, 4C overnight) and actin (Abcam ab49900) (1:10,000, 30 min RT).

### RNA isolation and RNA sequencing

A 6-well of organoids was collected in 800uL Trizol (Invitrogen, 15596-026) and immediately processed or stored at −80C. RNA was extracted according to the manufacturer’s instructions. DNA contamination was removed through treatment with recombinant DNaseI (Roche Diagnostics, #04716728001) for 15 minutes at RT and column purification using Qiagen RNeasy Mini kit (#74106). cDNA was prepared from 1ug of RNA (quantified by NanoDrop, Thermofisher Scientific Inc). Weill Cornell Medicine’s Genomics Core Laboratory checked RNA quality using a 2100 Bioanlyzer (Agilent Technologies), prepared the RNA library (TruSeq Stranded mRNA Sample Library Preparation kit (Illumina), and performed RNA sequencing (single end 75 cycles on a Illumina NextSeq 500). Raw NextSeq fastq files have been deposited in the sequence read archive (SRA) under accession PRJNA859154.

### RNAseq analysis

Raw FASTQ files were mapped to mouse (GRCm39) or human (GRCh38) reference genomes using STAR (v2.4.1d; default parameters)^49^. STAR count data was used for estimating differential gene expression using DESeq2^50^. For data visualization and gene ranking, log fold changes were adjusted using lfcShrink in DESeq2. R (v3.6.1) and R Studio (v1.2.1335) was used to create all visualizations and principal component analysis. Volcano plots and other visualizations were produced using the software packages:

Enhanced Volcano (https://bioconductor.org/packages/devel/bioc/html/EnhancedVolcano.html)
ggplot2 (https://cran.r-project.org/web/packages/ggplot2/index.html)

Variant calling was performed using picard (https://broadinstitute.github.io/picard/) and GATK (https://gatk.broadinstitute.org/hc/en-us) tools. Annotated SNPs in the mouse dbsnp database (ftp.ncbi.nlm.nih.gov/snp/organisms/mouse_10090/VCF/) were filtered from the analysis. The computational pipeline for picard and GATK, and code for processing variant tables and plotting is available at https://github.com/lukedow/iBE.git.

### Immunohistochemistry and immunofluorescence

Tissue was fixed, processed, and imaged as previously described^35^. IDEXX RADIL performed H&E on paraffin embedded sections. For immunofluorescence (IF), primary antibodies used were: rabbit anti-Cas9 (CST, #19526), mouse anti-p53 (CST, #2524), mouse anti-Glutamine synthetase (GS; BD Transduction Labs #610517), mouse anti-β-catenin (CST, #2698), rabbit anti-Cytokeratin-19 (CK19, Abcam, #ab133496), and rabbit anti-alpha Smooth muscle actin (aSMA, Abcam, #ab5694). Secondary antibodies used were donkey anti-rabbit 594 (1:500, Invitrogen, #A21207) and donkey anti-mouse 647 (Invitrogen, #A31571). All IF sections were counterstained with DAPI.

### Hydrodynamic teil vein injections

1ug SB13 transposase, 5ug SB-Myc, and gRNA (20ug plasmid gRNA or 2nmol Synthego synthetic standard chemically modified or 2nmol Synthego synthetic heavily modified gRNA51) in 2mL saline was delivered by lateral teil vein injection over 5-7s in 8-12 week old mice. Tumors were harvested after palpation and at a humane endpoint.

### Pancreas electroporation

Surgery to perform in vivo electroporation is previously described^52,53^. In brief, the surgical site is scrubbed with a povidone-iodine scrub (e.g., Betadine^®^, Nolvasan^®^), and the site is then rinsed with 70% alcohol. Under isofluorane (2-3%) anesthetization, a small laparotomy is performed and the pancreas is luxated with a blunt forceps. 5ug SB13 transposase, 25ug SB-KrasG12D, and gRNA (20ug gRNA plasmid or 2nmol Synthego synthetic, heavily modified gRNA51) in 30uL total volume (saline used to normalize) were delivered by injection into the pancreas. Solution is injected using a 27.5 gauge needle and tweezer electrodes are tightly placed around the injection bubble. Two pulses of electrical current using an in vivo electroporator (NEPAGENE NEPA21 Type II in vitro and in vivo electroporator) are applied. After electroporation, the peritoneum cavity is rinsed with 0.5ml of pre-warmed saline. Subsequently the peritoneum and muscles are sutured with absorbable sutures and the skin is closed with skin staples. The mice are kept at 37C until they are awake and post surgery pain management is done with injections of buprenorphine for the three following days (twice daily). Surgery and electroporation were performed on 8-12 week old mice. Tumors were harvested after palpation and at a humane endpoint.

### Statistical Analysis

All statistical tests used are indicated in the corresponding figure legends. In general, to compare two conditions, a standard two-tailed unpaired t test was used, assuming variance between samples. In most cases, analyses were performed with one-way or two-way ANOVA, with Tukey’s correction for multiple comparisons. Unless otherwise stated, each replicate represents an independent mouse/organoid lines or tumors from n ≥ 3 mice. Experimenters were not blinded to conditions. All statistics are reported in Supplementary Table 5.

---

**Supplementary Figure 1.**
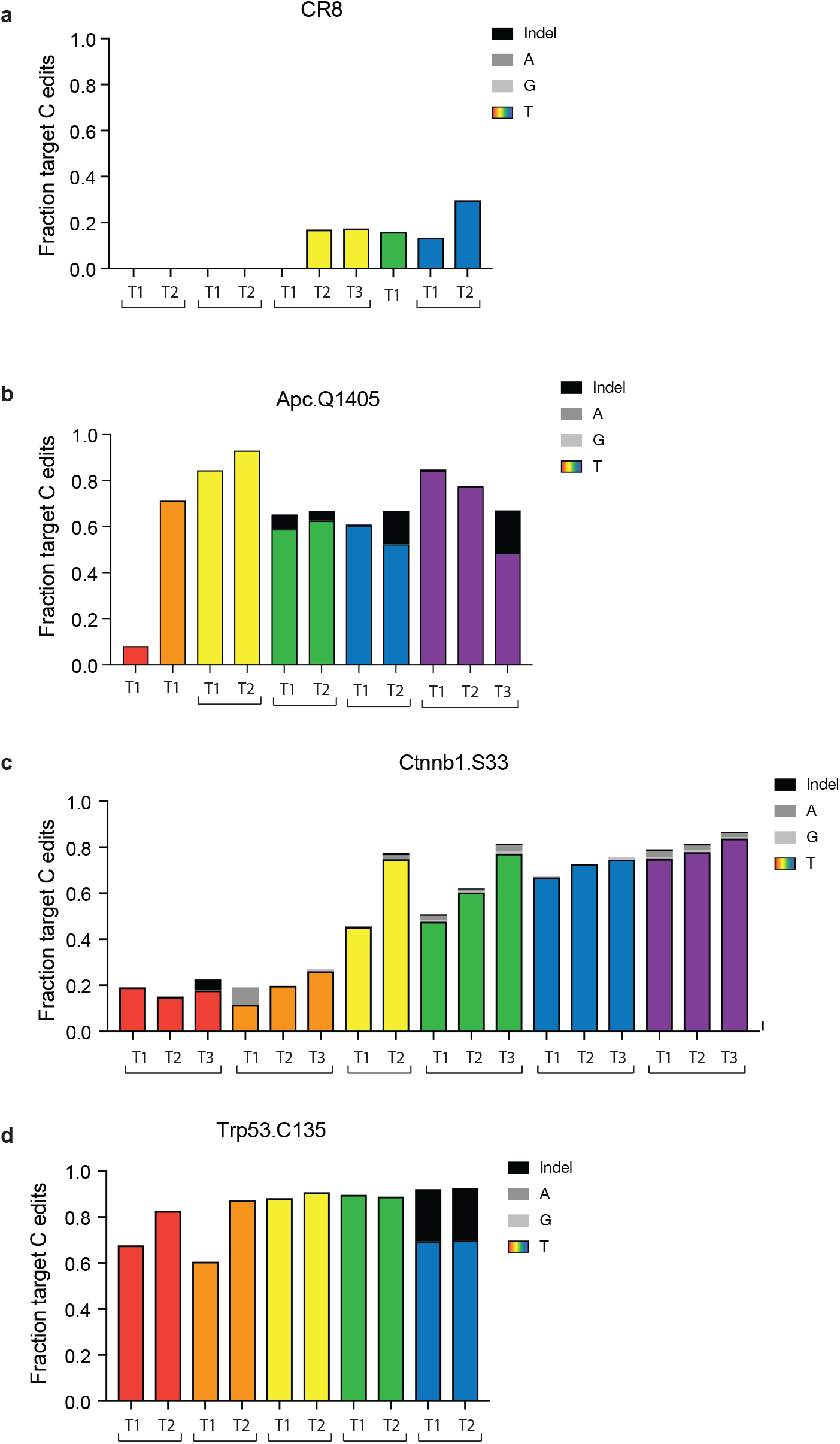
In situ base editing with iBE drives liver tumors. Targeted deep sequencing analysis of target C> T or other (A/G) edits or indels conversion for individual tumors corresponding to Figure 2c grouped by mouse. Each bar corresponds to an isolated bulk tumor with gRNA target listed above. a. CR8 b.Trp53Q135 c. ApcQ1405, and d. Ctnnb1S33.

**Supplementary Figure 2.**
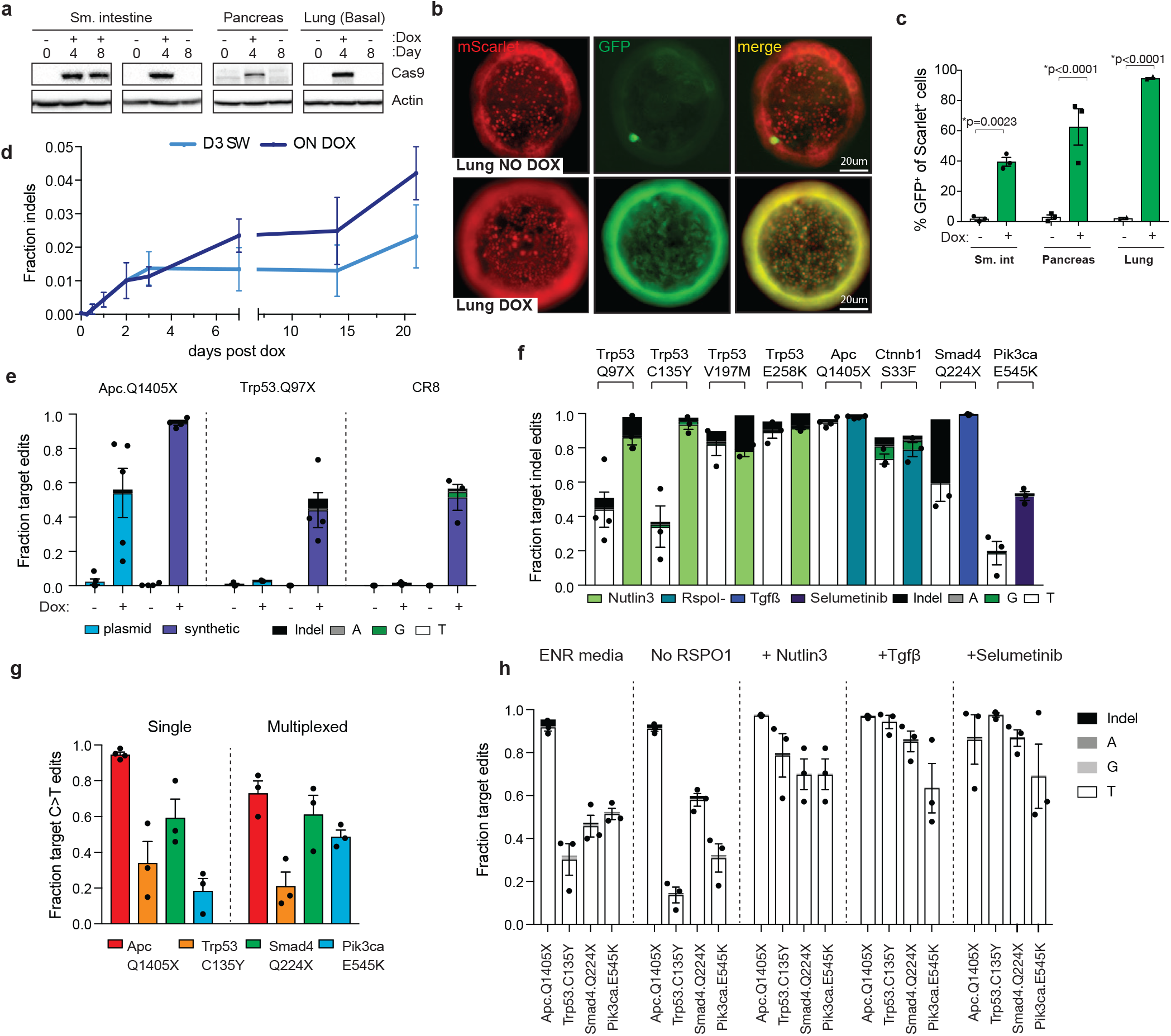
Efficient single and multiplexed base editing in iBE derived organoids. a. Cas9 immunoblot from organoids of each type (small intestine, pancreas, basal cells isolated from the mouse airway) cultured in dox naïve (Day 0, -dox), dox (Day 4, + dox), dox sustained (Day 8, + dox) or dox switched (Day8, -dox) conditions (protein taken as organoids undergo sequential dox conditions) in culture. ß-actin, loading control. b Fluorescence-based imaging of basal cell (lung) iBE organoids containing stable integration of GFPGO lentiviral construct cultured without (no dox) or with dox. Constitutive mScarlet marker for the reporter is shown in red (left column), base editing activatable GFP is shown in green (middle column), Merged images are depicted on the right. c. Quantification of GFPGO activation in mScarlet+ organoids with and without dox by flow cytometry. d. Targeted deep sequencing quantification of indel conversion at the ApcQ1405X locus in 2D small intestinal derived iBE cell line (constitutive shApc;shTrp53;KrasG12D cDNA) with lentiviral integration of a gRNA (LRT2B) targeting ApcQ1405 at time points after dox is added to the culture media (0 to 15 days) (dark blue),or dox was cultured in the media briefly for 3 days (light blue). e. Targeted deep sequencing quantification of corresponding target C>T/A/G and indel conversion in small intestinal iBE organoids nucleofected with various plasmid (light blue) or synthetic (indigo) gRNAs (ApcQ1405, Trp53Q97, CR8.OS2) with and without dox conditions for 2 days surrounding nucleofection. f. Targeted deep sequencing quantification of target C>T/A/G and indel conversion in small intestinal iBE organoids nucleofected with various synthetic gRNAs targeting cancer associated SNVs cultured in dox or 2 days surrounding nucleofection, and either unselected (-) or selected with corresponding functional selective media condition (+: Trp53mut, nutlin selection; Smad4mut, Tgfß selection; Pik3caact, selumetinib selection; Ctnnb1act, RspoI withdrawal). g. Targeted deep sequencing of C>T editing at target sites in single targeted versus quadruple targeted (multiplexed) organoids at each corresponding site (ApcQ1405, red; Trp53C135,orange; Smad4Q224, green;, Pik3caE545, blue) h. Targeted deep sequencing quantification of target C>T/A/G and indel conversion (and C> other or indels) in small intestinal iBE organoids nucleofected with 4 synthetic gRNAs in f (green boxes) at each gRNA target loci (ApcQ1405, Trp53C135, Smad4Q224, Pik3caE545). Media conditions and corresponding organoid genotype and sequencing information is grouped and listed above.

**Supplementary Figure 3.**
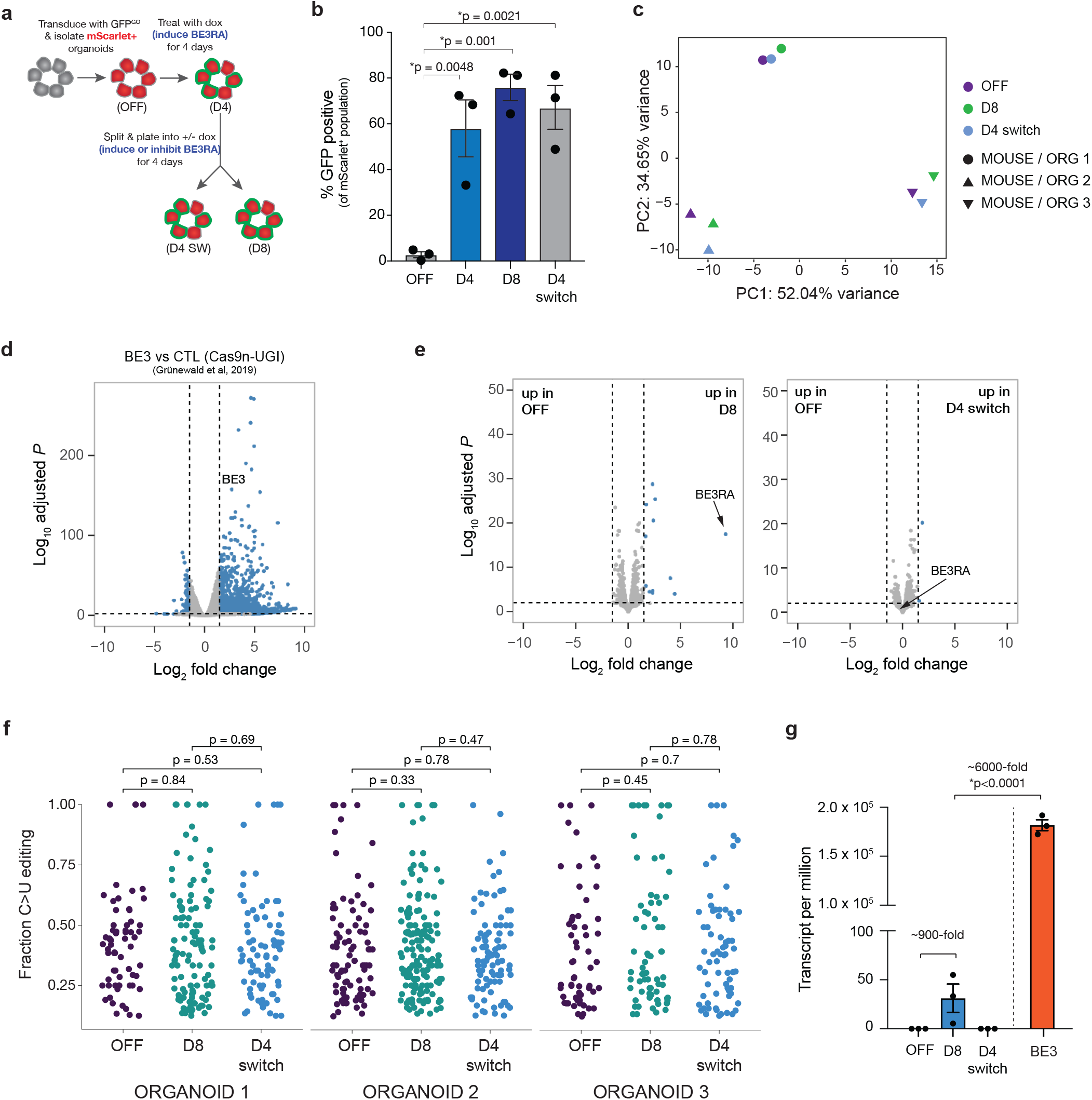
iBE does not induce off target RNA editing. a. Schematic of experimental set up in iBE derived pancreatic organoids. Organoids were transduced with GFPGO construct and selected for reporter containing organoids (mScarlet+). Organoids maintained off dox were then split into dox conditions to induce CBE expression for 4 days and then split again into + and – dox conditions for an additional days. b. Organoids at each conditions labeled (OFF, D4, D8, and D4 sw) were quantified by flow cytometry for DNA on target activity reported by BE activated GFP expression. GFP+ cells are quantified within the mScarlet+ population. c. PCA analysis of RNA isolated samples from OFF, D8, and D4 sw organoid conditions. Colors correspond to dox condition and shape delineates organoid replicate/mouse origin (n=3). d. Differential gene expression analysis of D8 vs. OFF (left) and D4 sw vs. OFF (right). e. Differential gene expression analysis of transfection-based BE3 overexpression in HEK293T cells compared to control Cas9n-UGI overexpressed cells (without rAPOBEC1) f. Relative transcript abundance (per million) of OFF, D8 and D4 sw organoid samples. BE3 transfected HEK293Ts plotted on right for indirect comparison. Fold changes are described between BE3 induced conditions. g. C to U editing in RNA transcripts detected from RNA sequencing data from OFF, D8, and D4 switched organoids. RNA editing from three independent organoids replicates are shown.

**Supplementary Figure 4.**
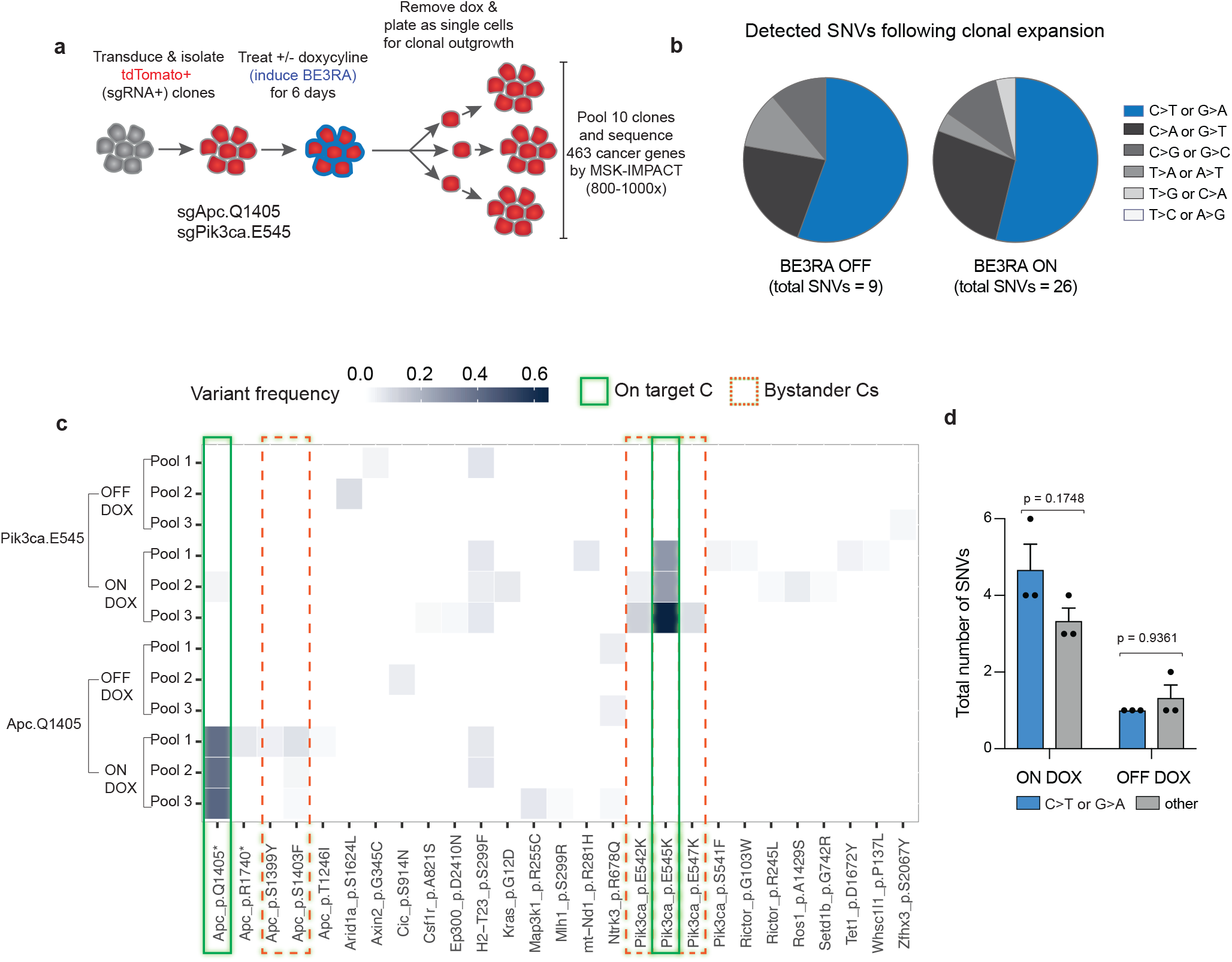
iBE has low DNA off target activity. **a.** Schematic of experimental set up in mouse embryonic stem cells (ESCs). mESCs containing iBE knock in were transduced with LRT2B-gRNA vector and selected for gRNA expression. sgRNA+ cells were plated with and without dox for 6 days after which cells were plated at low density for clonal outgrowth without dox. 3 pools of 10 clones were picked for each dox conditions across to gRNA targeted cell lines (gRNAs = Apc1405 and Pik3caE545). Clones were sent for deep sequencing across the MSK-IMPACT cancer gene set for 800-100X read coverage. b. Pie chart display of frequency of C to T or C to other SNVs found in pooled clones for each condition (on and off dox) for both gRNAs. b. Sequencing analysis at cancer gene sites in cell conditions (right) described in a. Solid green boxes are on target activity of gRNA, Dotted green boxes signify on target bystander editing within the gRNA window. d. Quantification of C to T and C to other SNVs found across both targets. 2-way ANOVA test for multiple comparisons was used to evaluate statistical significance across conditions. p-values are displayed.

**Supplementary Figure 5.**
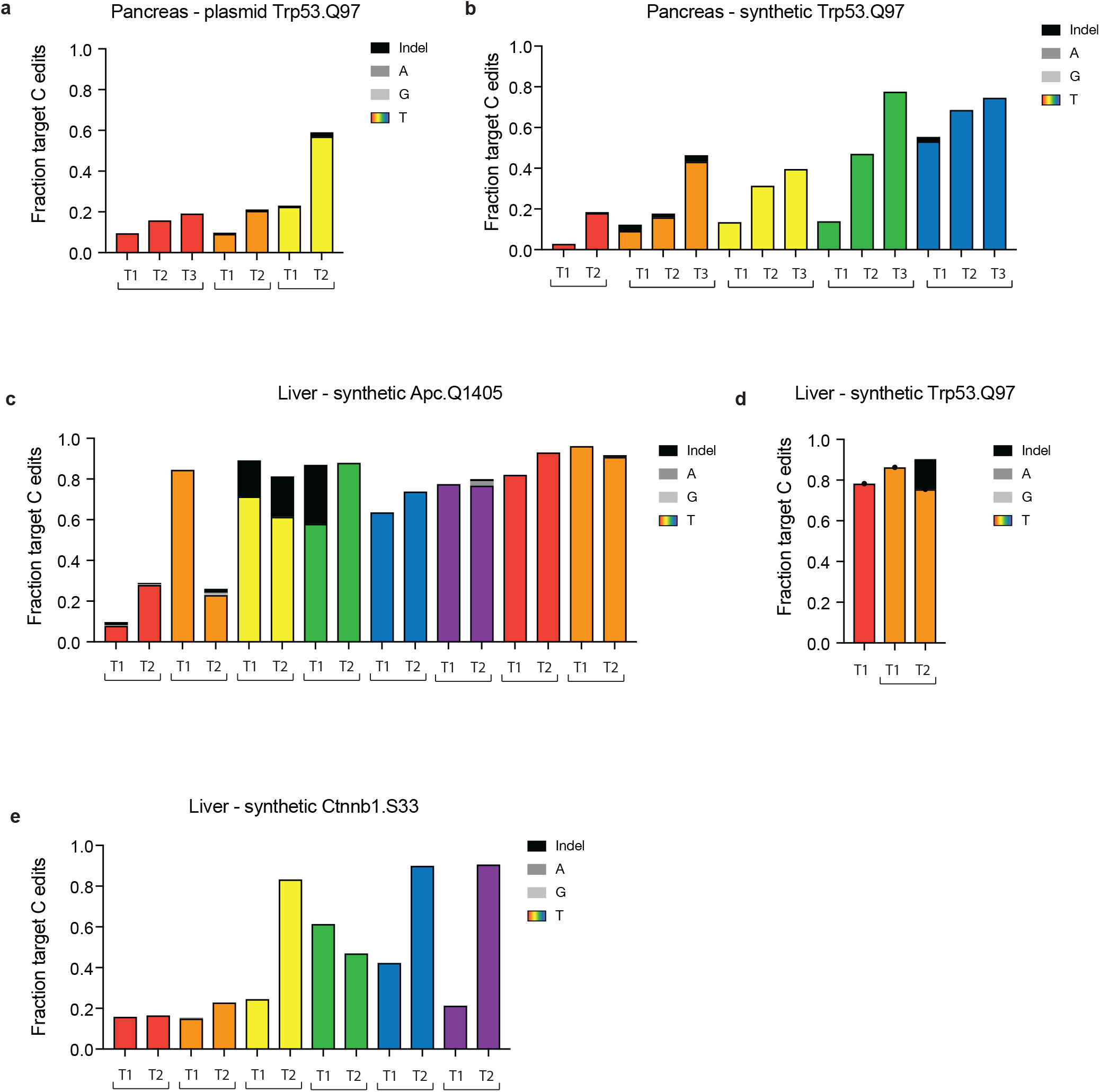
In situ base editing with iBE drives liver tumors. Targeted deep sequencing analysis of target C> T or other (A/G) edits or indels in individual tumors corresponding to Figure 4c and d grouped by mouse. Each bar corresponds to an isolated bulk tumor with tumor type, gRNA type and target listed above. a. Trp53Q97X targeted with plasmid guide (LRT2B) in pancreas, b.Trp53Q97X targeted with synthetic guide in pancreas c. ApcQ1405 targeted with synthetic guide in liver d. Trp53Q97X targeted with synthetic guide in liver, e, Ctnnb1S33 targeted with synthetic guide in liver.

**Supplementary Figure 6.**
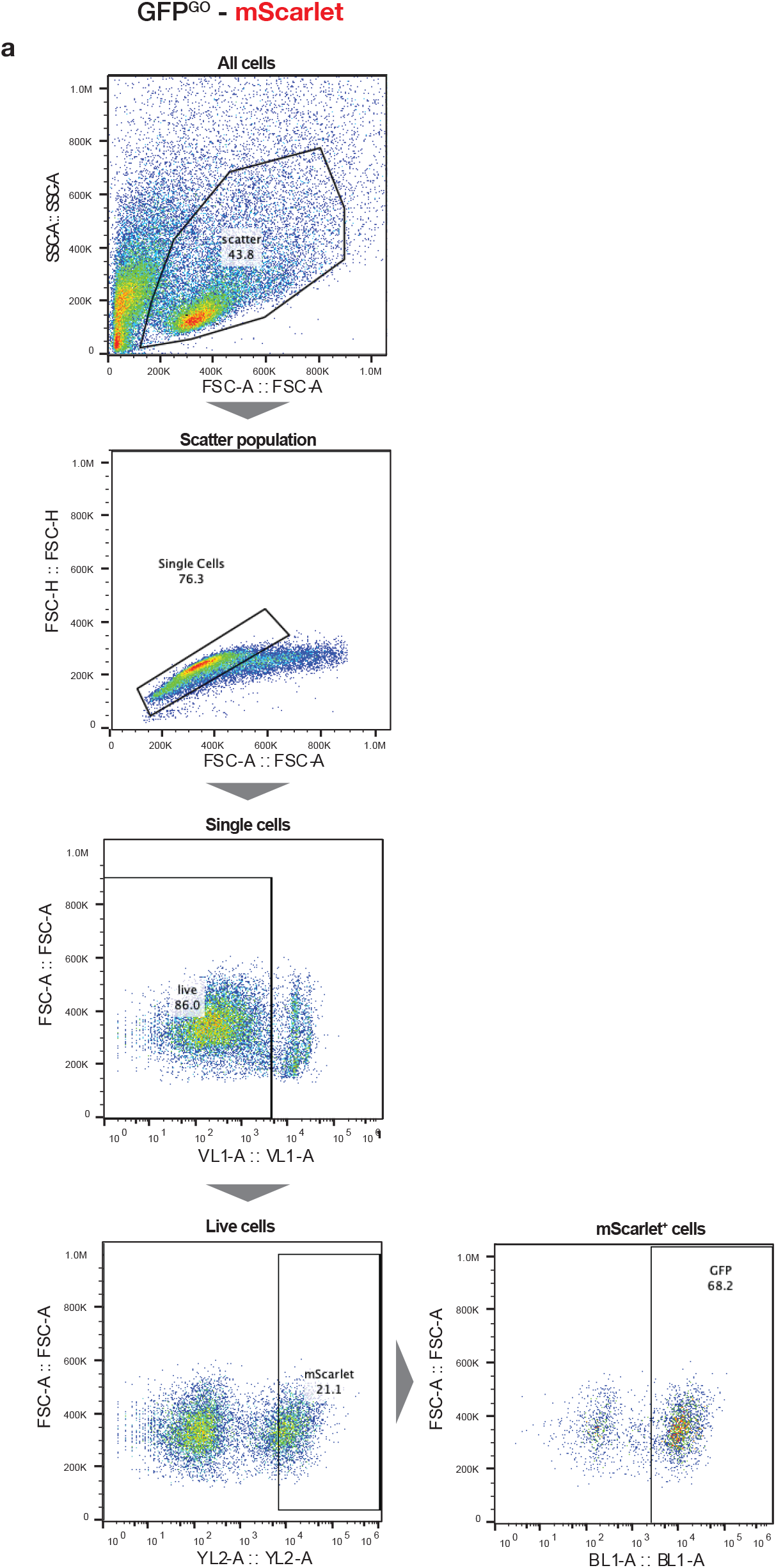
Representative gating for flow cytometry analysis of editing with GO reporter. From top to bottom: stepwise gating of live, single, DAPI-negative cells, mScarlet positive cells for quantification of GFP percentage (lower right)

